# Temporal Microenvironment Mapping (μMap) of Intracellular Trafficking Pathways of Cell-Penetrating Peptides Across the Blood-Brain Barrier

**DOI:** 10.1101/2025.01.15.633151

**Authors:** Danielle C. Morgan, Steve D. Knutson, Chenmengxiao (Roderick) Pan, David W. C. MacMillan

## Abstract

Peptides play critical roles in cellular functions such as signaling and immune regulation, and peptide-based biotherapeutics show great promise for treating various diseases. Among these, cell-penetrating peptides (CPPs) are particularly valuable for drug delivery due to their ability to cross cell membranes. However, the mechanisms underlying CPP-mediated transport, especially across the blood-brain barrier (BBB), remain poorly understood. Mapping intracellular CPP pathways is essential for advancing drug delivery systems, particularly for neurological disorders, as understanding how CPPs navigate the complex environment of the BBB could enable the development of more effective brain-targeted therapies. Here, we leverage a nanoscale proximity labeling technique, termed µMap, to precisely probe the peptide– receptor interactions and intracellular trafficking mechanisms of photocatalyst-tagged CPPs. The unique advantage of the μMap platform lies in the ability to control the timing of light exposure, which enables the collection of time-gated data, depending on when the blue light is applied to the cells. By harnessing this spatiotemporal precision, we can uncover key peptide–receptor interactions and cellular processes, setting the stage for new innovations in drug design and brain-targeted therapies.

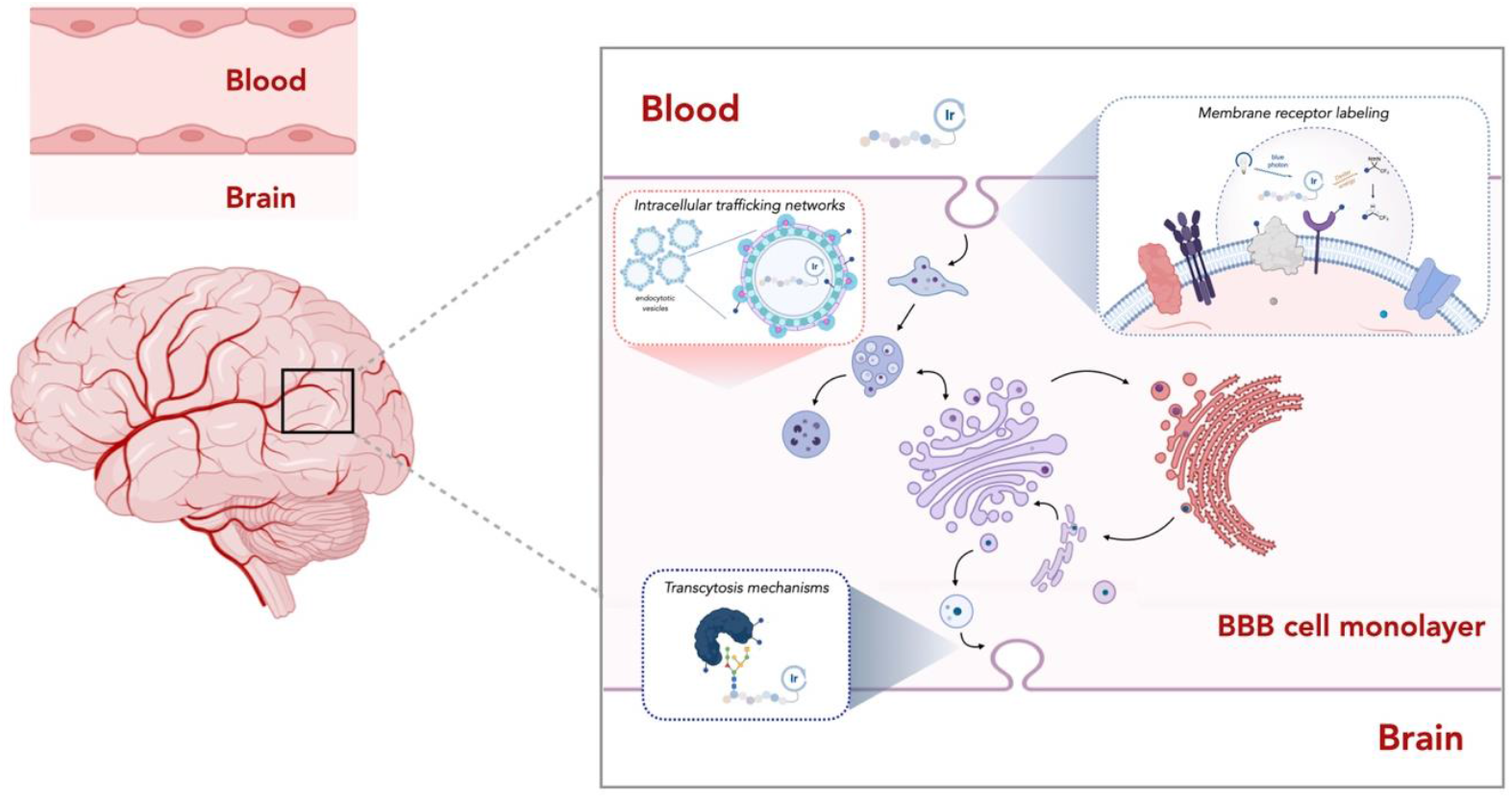

## Introduction

The effective delivery of small molecule drugs and biologics, particularly across challenging barriers such as the blood-brain barrier (BBB), offers substantial promise for therapeutic intervention in a wide range of neurological diseases.^[1-3]^ However, the size and hydrophilic nature of these molecules often limits their ability to cross cell membranes and restricts their therapeutic potential.^[4]^ Cell-penetrating peptides (CPPs) have emerged as a promising solution to drive transport of diverse molecules, including large, biologically active proteins, into cells through endocytosis and across cell membranes via transcytosis.^[5-7]^

Most CPPs are positively charged at physiological pH due to the presence of arginine and/or lysine residues, and it is well established that these peptides interact with negatively charged proteoglycans on the cell surface to facilitate endocytic internalization.^[8-11]^ However, a subset of CPPs that lack positive charge also exhibit cell-penetrating capabilities, raising key questions about the underlying mechanisms of their membrane interaction, translocation, and cellular entry.^[12-14]^

Moreover, many CPPs possess the ability to not only enter cells but also exit through a process known as transcytosis.^[15, 16]^ These specialized peptides can traverse entire cell monolayer barriers such as the BBB and thus hold promise as drug delivery systems to treat neurological conditions that are difficult to address with traditional methods.^[17]^ Profiling transcytosis pathways at the BBB is crucial for understanding how molecules move from the bloodstream into the brain.^[1]^ Transcytosis, the transport of molecules across endothelial cells, plays a key role in how substances like drugs, antibodies, and hormones cross the BBB.^[18]^ By unraveling the mechanisms behind these transcytosis pathways, we can identify strategies to enhance or bypass the BBB for more effective drug delivery, particularly for neurological diseases like Alzheimer’s, Parkinson’s, and multiple sclerosis, where the BBB may be compromised or overly restrictive.^[19]^ Identifying key transporters and receptors could enable the development of targeted therapies that facilitate drug delivery to the brain. However, the precise mechanisms by which CPPs engage in cellular entry and transcytosis remain poorly understood, and a deeper understanding of these processes could significantly accelerate the development of effective drug delivery strategies, particularly for the targeted treatment of brain disorders.

To this end, our laboratory has introduced a photocatalytic proximity labelling technology termed µMap, which enables precise labeling of cell-surface protein–protein interactions with exceptional spatial resolution.^[20-26]^ More recently, we have broadened the capabilities of µMap to investigate intracellular systems through photocatalyst design and cellular engineering approaches.^[24, 26, 27]^ The intracellular µMap platform offers significant advantages over traditional proximity labeling (PL) methods, such as BioID and APEX, by providing high-resolution temporal control and minimizing cellular disruption.^[28, 29]^ These features are crucial when studying the dynamic and complex processes involved in cellular trafficking, where rapid labeling and minimal interference with cellular function are essential. ^[21, 22, 26]^

We recognized that the spatiotemporal precision of µMap could make it a powerful tool for studying the trafficking dynamics of cellular delivery mechanisms, such as those involving CPPs. Here in, we describe the development of a µMap-based technology to elucidate the intracellular trafficking pathways of photocatalyst-tagged CPPs. Crucially, this platform can be tailored with short CPP-iridium incubation times to selectively enrich potential cell surface entry receptors, or longer incubation periods to uncover cellular trafficking pathways and transcytosis mechanisms (Figure 1).

**Fig 1.**
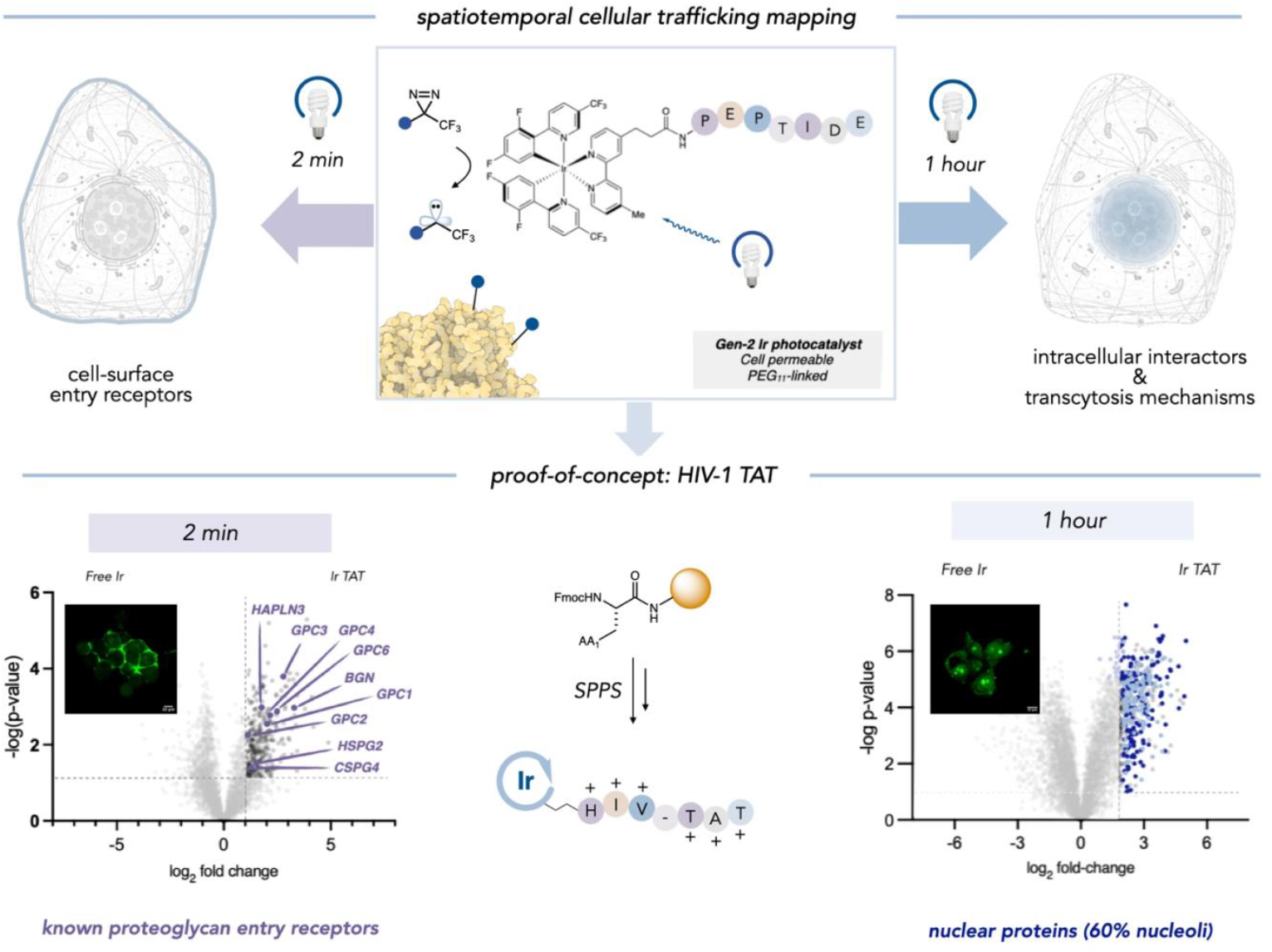
The peptide μMap platform. The spatiotemporal control of μMap enables time-resolved labeling of cellular trafficking events. HIV-1 TAT was used to validate the platform, capturing cell-surface entry receptors at 2 minutes and intracellular interactors at 1 hour, thus providing insights into dynamic, time-dependent trafficking processes.

We first sought to validate the platform in the context of a well-studied CPP, HIV-1 TAT, which is known to have electrostatic interactions between its basic amino acids and negatively charged proteoglycans on the cell surface.^[10]^ HIV-1 TAT is widely recognized for its ability to facilitate the transmembrane delivery of various therapeutic cargoes, including small molecules,^[30]^ peptides/proteins,^[31, 32]^ antibodies,^[33]^ liposomes,^[34]^ nanoparticles,^[35]^ small interfering RNAs (siRNA)^[36]^ and antisense oligonucleotides.^[37, 38]^ Many of these cargoes have been investigated in clinical trials for their potential therapeutic applications. Our primary focus, however, was to assess the platform’s capability with CPPs whose entry receptors were previously unknown. Using μMap cell-surface labeling experiments, we successfully identified unique entry receptors for two distinct CPPs. Building on these insights, our ultimate objective was to harness the μMap platform to profile the transcytosis pathways across the BBB.

Overall, we focused on selecting CPPs with unknown cellular trafficking mechanisms, including one that demonstrates the ability to efficiently cross the BBB and selectively target specific brain regions.^[39]^ Iridium-peptide conjugates were synthesized for intracellular mapping, enabling the identification of cellular trafficking partners and previously unknown entry receptors for all the selected peptides.

## Results and discussion

### Proof-of-Concept: HIV-TAT

As a proof-of-concept of the μMap-ID platform, we first sought to evaluate the endocytosis and intracellular trafficking of a well-characterized CPP, HIV-TAT.^[8-12]^ Before conducting μMap proximity labeling, we first assessed the internalization kinetics of a fluorescein-labeled HIV-1 TAT peptide in HEK293T cells via fluorescence microscopy (Figure 2A). We observed that, over a time course of 2 minutes to 1 hour, the HIV-1 TAT peptide is rapidly internalized into cells and localizes in the nucleus/nucleoli. With this information in hand, we generated the HIV-1 TAT peptide conjugated to a cell-permeable iridium photocatalyst via solid-phase peptide synthesis (SPPS). Notably, the photocatalyst is compatible with the SPPS workflow, including the acidic conditions required for peptide cleavage from solid support. The cytotoxicity of the peptide-iridium conjugate was evaluated at timepoints and concentrations that mirrored microscopy and subsequent labeling parameters (2 µM, 2 min to 1 hour). As shown in Figure 2C, relatively low toxicity was observed with the iridium-TAT conjugate, as well as with a free-iridium control.

**Fig 2.**
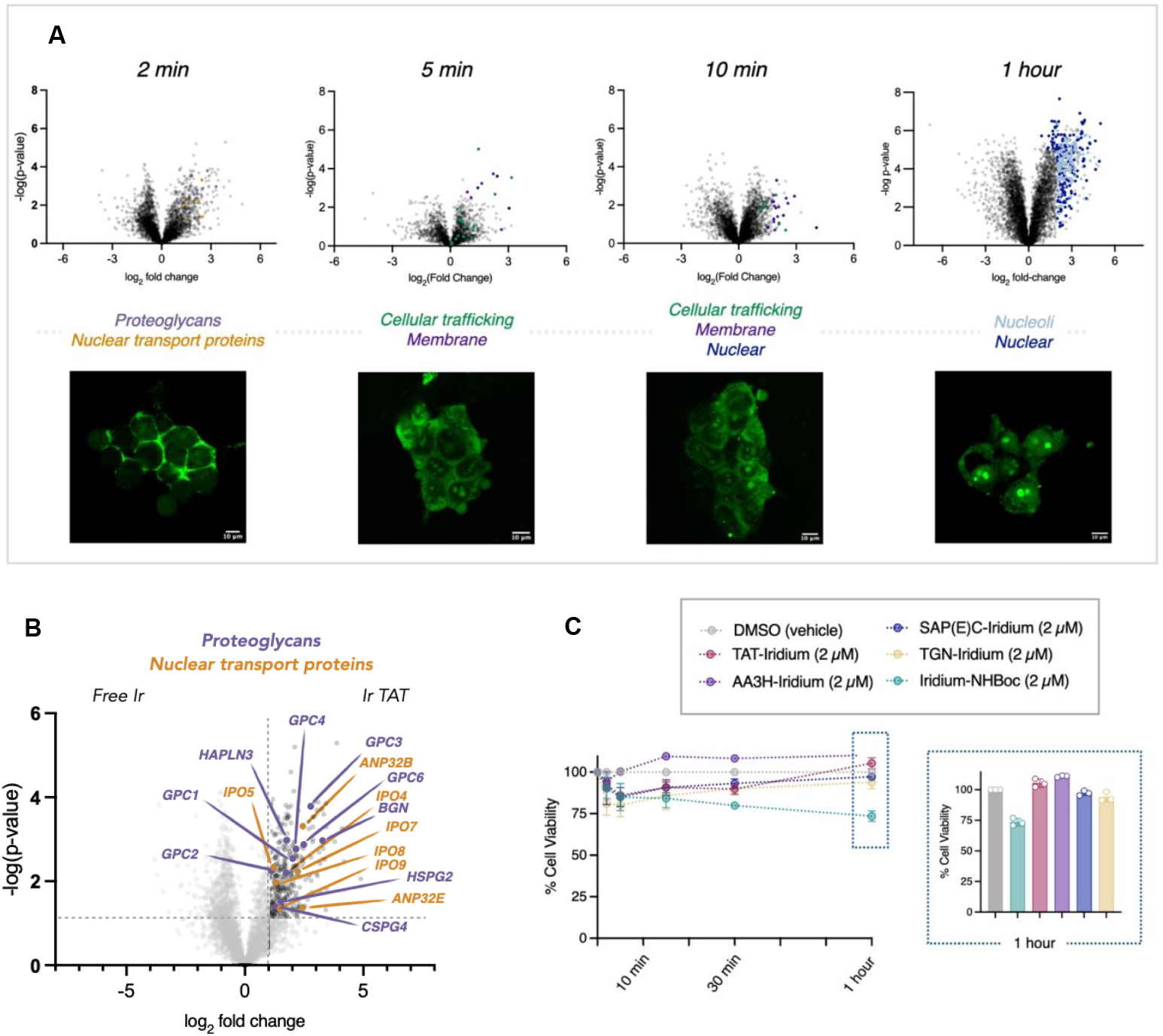
Spatiotemporal profiling of HIV-1 TAT subcellular trafficking. **A.** Parallel microscopy and μMap analysis were employed to examine the intracellular trafficking of the Iridium-HIV-1 TAT peptide, providing both visual and proteomic insights. **B**. Early interactome analysis identified cell surface receptors and proteoglycans (*HSPG2, CSPG4, GPC1–4, GPC6, BGN, HAPLN3*, purple) known to facilitate internalization. Additionally, nuclear import factors (importins *IPO4, IPO5, IPO7, IPO8, IPO9*, yellow) were detected. **C**. Toxicity assessments were conducted on all peptide-Iridium conjugates used in this study, evaluating their effects at a concentration of 2 μM over the maximum incubation time (1 hour) in cells.

Based on the initial studies, we selected “early” (2 minutes), “middle” (5/10 minutes) and “late” (1 hour) time points for the labeling experiments. Cells on ice were exposed to Ir-TAT for the selected time points, then washed with cold Dulbecco’s Phosphate Buffered Saline (DPBS) and treated with diazirine-PEG3-biotin. Subsequently, the cells were irradiated with blue light for three minutes to label interacting protein partners of HIV-1 TAT based on their proximity to the peptide. Proteomic analysis of the Ir-TAT interactome at the 2-minute mark (Figure 2B) revealed several cell surface proteins, including the proteoglycans *HSPG2, CSPG4, GPC1-4, GPC6, BGN* and *HAPLN3*, which have been previously shown to facilitate the internalization process of cationic CPPs.^[9-11]^ Additionally, at the same time point (2 min) we identified nuclear import proteins (importins; IPO4, IPO5, IPO7, IPO8, IPO9), which have previously been reported to be involved in the nuclear translocation of HIV-TAT.^[40, 41]^ µMap profiling at the 10-minute time point revealed several known trafficking mediators, such as Ras-related proteins (RABs), indicating vesicular localization occurring as part of endocytosis. Longer exposure labeling experiments (1 hour) displayed 82% of enriched hits as nuclear proteins (60% nucleoli), mirroring our initial microscopy observations (Figure 2A). Overall, these initial experiments successfully traced HIV-1 TAT subcellular trafficking within cells in a temporal manner dependent upon the peptide-iridium incubation time.

### Evaluating unknown entry mechanisms of non-cationic CPPs

Building on these results, we next sought to investigate CPPs with less well-understood entry mechanisms, particularly those that do not possess the characteristic positive charges. As CPPs of interest, we identified the neutral peptide AA3H and the negatively charged peptide SAP(E)C (Figure 3A), both of which have been hypothesized to enter cells via active transport, though their specific entry receptor(s) remain unclear.[42, 43]

**Fig 3.**
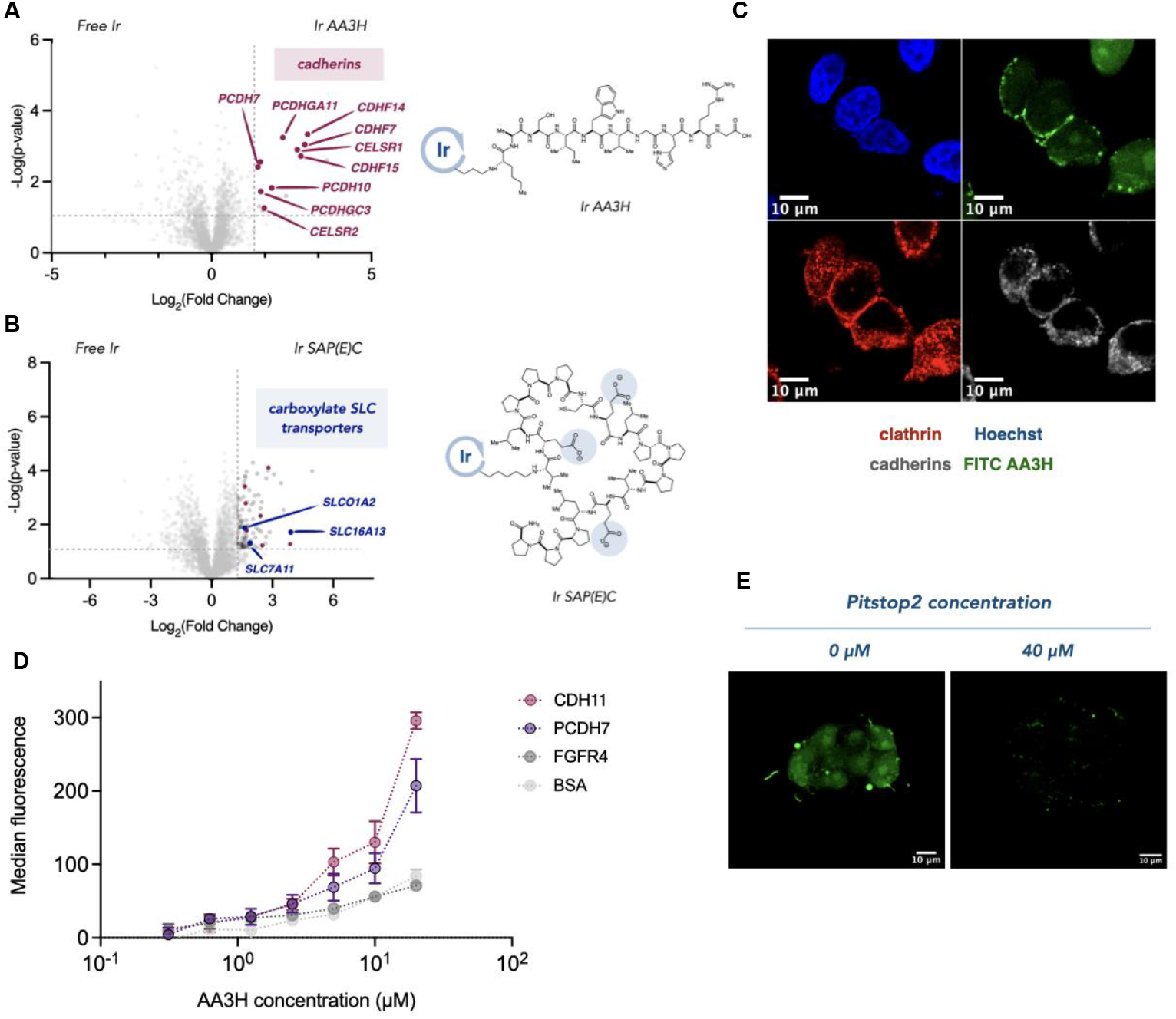
Identifying the entry receptors for neutral AA3H and anionic SAP(E)C CPPs. **A.** μMap proteomic analysis of AA3H entry identified cadherins (red) and **B**. SLC transporters (blue) as interactors for SAP(E)C. **C**. Immunofluorescence microscopy displays colocalization of FITC-AA3H with cadherins and clathrin-coated pits (scale bar = 10 μm). **D**. ELISA binding studies demonstrate significant interaction between AA3H and cadherins, with minimal binding to control proteins. **E**. Microscopic evaluation of AA3H entry is disrupted *via* inhibition of clathrin-mediated endocytosis (scale bar = 10 μm).

We performed experiments analogous to those conducted with HIV-1 TAT peptides, starting with fluorescence microscopy to determine optimal time points for internalization. Notably, both peptides demonstrated slower cellular uptake, with fluorescein-labeled peptides becoming visible inside cells only after 1 hour. Based on these observations, μMap experiments were carried out by incubating HEK293T cells with iridium-conjugated AA3H or SAP(E)C peptides for 1 hour, followed by blue light irradiation.

Our results revealed a clear distinction between enriched hits associated with the neutral and negatively charged peptides (Figure 3A). µMap profiling of the negatively charged SAP(E)C peptide revealed an enrichment of several carboxylate SLC transporter proteins, including SLCO1A1. Notably, SLCO1A1 has previously been identified as a facilitator of anionic natural product peptide transport, suggesting it may function as a potential entry receptor for SAP(E)C.^[44]^

In contrast, the neutral AA3H peptide displayed striking enrichment of several cell-adhesion cadherin proteins. Interestingly, cadherins are internalized *via* clathrin-mediated endocytosis as part of their role in cell-cell adhesion, raising the possibility that AA3H may exploit this mechanism for internalization (Figure 3B). To investigate this hypothesis further, we performed microscopy to examine the localization of cadherins, clathrin pits, and AA3H (Figure 3C). Notably, we observed that treatment of cells with an inhibitor of clathrin-mediated endocytosis, Pitstop2 (40 μM), disrupted AA3H internalization, offering additional evidence for a clathrin-dependent uptake mechanism (Figure 3E). To directly assess the interaction between AA3H and cadherins, we conducted an Enzyme-Linked Immunosorbent Assay (ELISA) assay using immobilized cadherins (CDH11 and PCHD7) identified in our μMap experiments. We observed significant binding of AA3H to both cadherins and only minimal binding with control proteins BSA and FGFR4 (Figure 3D). Moreover, the negatively charged SAP(E)C peptide showed negligible binding to these cadherins, further supporting the specificity of the interaction between AA3H and cadherins.

### Exploring transcytosis mechanisms of blood-brain barrier (BBB) shuttle peptides

We next sought to leverage the µMap platform for a rigorous investigation into the mechanisms of BBB transcytosis. For this study, we identified PepTGN, a linear peptide discovered through an *in vivo* phage display campaign that is capable of permeating the BBB *via* a cellular mechanism that remains to be elucidated.

### Identifying the entry receptor of PepTGN

A key advantage of the peptideID μMap platform is the ability to conduct live-cell experiments across a wide range of cell lines. Given the limited information available on the binding kinetics of PepTGN and its potential receptors, we selected two separate cell lines for the experiments: the mouse neuronal endothelial cell line bEND.3 and the widely used hCMEC/d3 human BBB model. To identify a potential cell-surface entry receptor for PepTGN, we first performed μMap experiments on ice to prevent any active transport-mediated internalization. Proteomic comparisons of both datasets showed enrichment of known membrane proteins (*SNX4, THBS1, CALCRL, AHSG* and *NECAP2*), which were prioritized for further investigation as potential entry receptors.

Specifically, the Calcitonin receptor-like receptor (*CALCRL*) was identified as an intriguing cell-surface receptor hit across both mouse and human BBB experiments (Figure 4B, human BBB experiment). *CALCRL* is a G-protein coupled receptor (GPCR) known to bind and internalize calcitonin peptide hormones *via* clathrin- and dynamin-dependent endocytosis. Interestingly, top enriched hits also included proteins associated with clathrin (*GAK* and *NECAP2*) and dynamin (*DNM2*), suggesting a similar endocytic pathway for PepTGN (Figure 4B). Upon sequence analysis, we observed intriguing similarities between PepTGN and the calcitonin peptide hormones, particularly in the C-terminal region. We thus hypothesized that PepTGN interacts with *CALCRL* at the cell surface and is subsequently internalized through receptor-mediated endocytosis.

**Fig 4.**
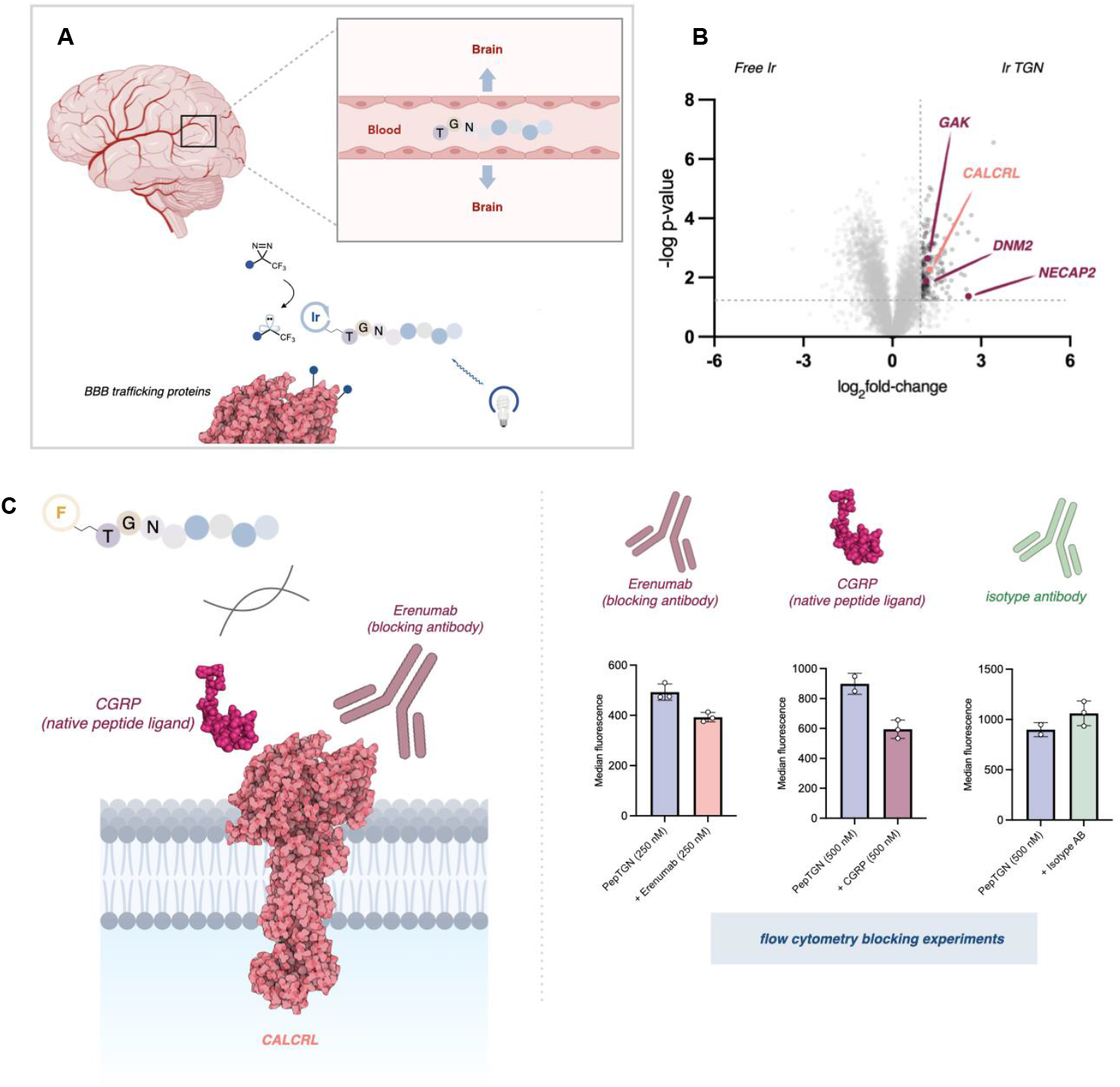
Entry receptor mapping of BBB penetrating PepTGN identified through *in vivo* HTS. **A.** PepTGN, a brain-penetrating peptide identified *via in vivo* phage display, has an unknown BBB entry mechanism. We utilized μMap to investigate its transcytosis trafficking pathway. **B**. μMap experiments were performed on ice for 15 minutes in two different cell lines (hCMEC/d3, shown, and bEND.3 cells) to identify potential entry receptors. One of the top hits identified in both volcano plots was the calcitonin receptor-like receptor (*CALCRL, pink*), a G-protein coupled receptor (GPCR) involved in peptide signaling. The experiments also revealed clathrin (*GAK* and *NECAP2*, maroon) and dynamin (*DNM2*, maroon) proteins. **C**. Blocking experiments were conducted to validate *CALCRL* as the entry receptor for PepTGN. These studies showed a significant reduction in peptide uptake when cells were pre-incubated with either the native ligand CGRP or the monoclonal antibody Erenumab. In contrast, incubation with a control isotype antibody had no effect on internalization.

To further investigate whether *CALCRL* serves as an entry receptor, we assessed internalization of a FITC-labeled PepTGN using flow cytometry in the presence of various blocking agents. Specifically, we used the native ligand calcitonin gene-related peptide (α-CGRP), as well as Erenumab (Aimovig), a monoclonal antibody used in migraine therapy that targets *CALCRL* and inhibits CGRP binding. PepTGN internalization was significantly decreased when cells were pre-incubated with Erenumab or the α-CGRP native ligand compared to a non-binding isotype control (Figure 4C), suggesting that *CALCRL* is involved in the endocytosis mechanism of PepTGN.

We believe that this demonstrates how the μMap platform can be used to identify BBB receptors that can be targeted and utilized for BBB drug delivery.

### Mapping the intracellular trafficking and transcytosis pathway of PepTGN for brain entry

Having identified a potential entry pathway for PepTGN, we next harnessed the spatiotemporal profiling capabilities of μMap to investigate its intracellular trafficking mechanisms.

While our experiments to identify cell-surface entry receptors were conducted at 4 ºC, we tracked the intracellular movement and subsequent exocytosis of PepTGN at higher temperatures (37 ºC) over a longer incubation time (1 h). As expected, top protein interactors from this experiment were predominantly intracellular (80%), with about 30% of these genes associated to endoplasmic reticulum (ER) and Golgi trafficking as well as secretory pathways (Figure 5A, bEND.3 cell line), suggesting a potential cellular route for exocytosis.

**Fig 5.**
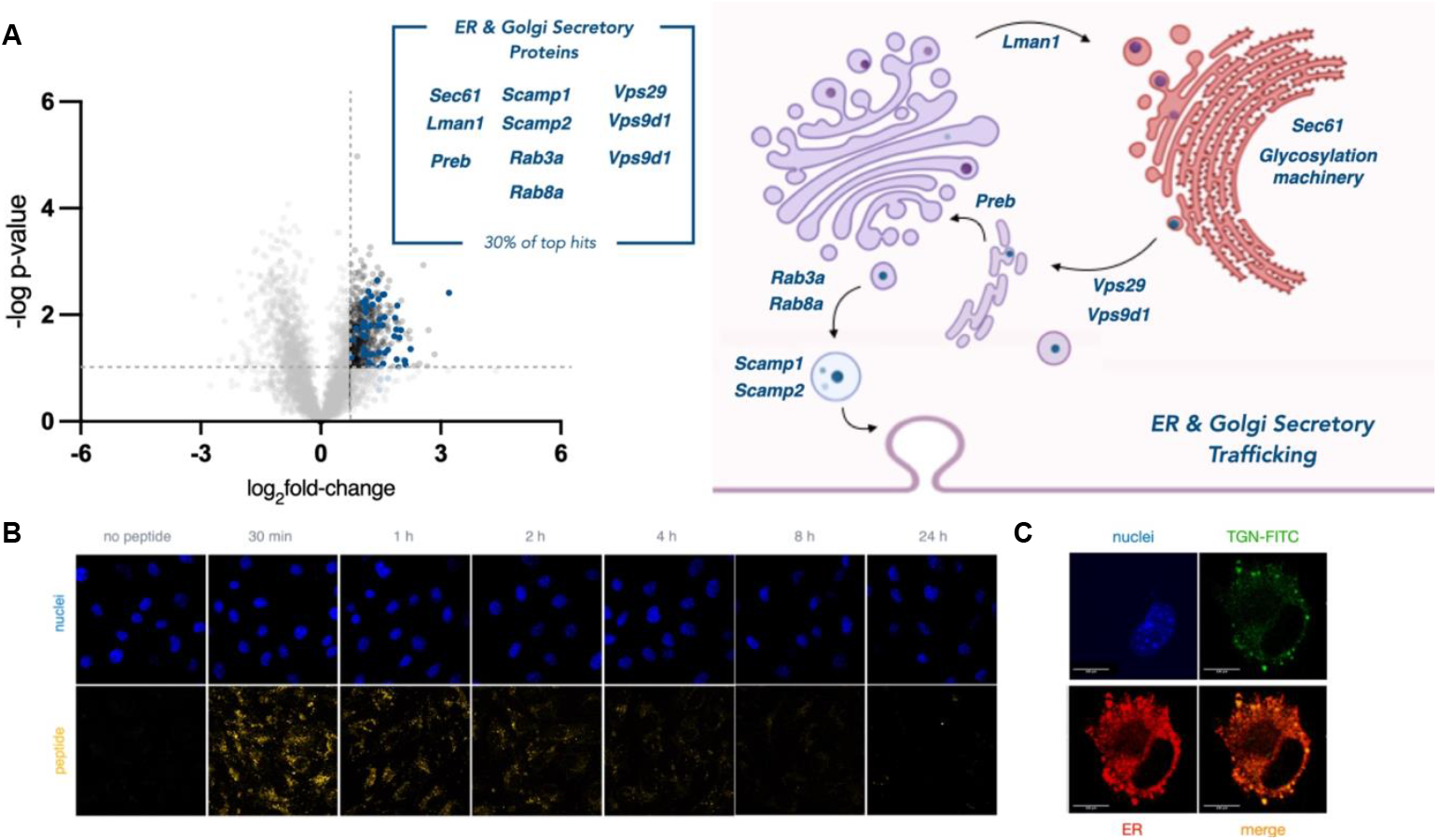
Transcytosis mapping of BBB penetrating PepTGN identified through *in vivo* HTS. **A.** μMap profiling of PepTGN intracellular interactors in bEND.3 cell line after 1 hour incubation. 30% of the top hits in the volcano plot relate to ER and Golgi secretory proteins (blue), suggesting a potential exocytosis mechanism of PepTGN. **B**. Fluorescence microscopy of secretion kinetics of FITC PepTGN. **C**. FITC-PepTGN (green) colocalization in the ER (red, ERp57/ERp60 antibody) (scale bar = 100 μm).

We next assessed secretion kinetics and cellular localization of PepTGN with immunofluorescence microscopy (Figure 5B,C). Notably, FITC-PepTGN was rapidly internalized (<30 minutes at 37 ºC) and undetectable after ∼4 h, indicating exocytosis within this time frame. PepTGN was predominantly localized within distinct puncta after 30 minutes incubation (Figure 5C), overlaying with an ER-marker (ERp57/ERp60 antibody) in agreement with interactors identified in μMap experiments (Figure 5A).

We also noted that O-glycosyltransferases (such as *Galnt12, Galnt2, Galnt4, Eogt, C1galt1c1*, and *Mlec*) were among the top hits in the bEND.3 cell line experiment, prompting us to investigate whether PepTGN undergoes glycosylation during secretion *via* the ER and Golgi. To explore this possibility, we first treated cells with fluorescein-labeled PepTGN and then isolated the secreted peptide from media. Upon analyzing this material *via* gel electrophoresis, we observed the emergence of two higher molecular weight bands, suggesting possible glycosylation events (Figure 6B).

**Fig 6.**
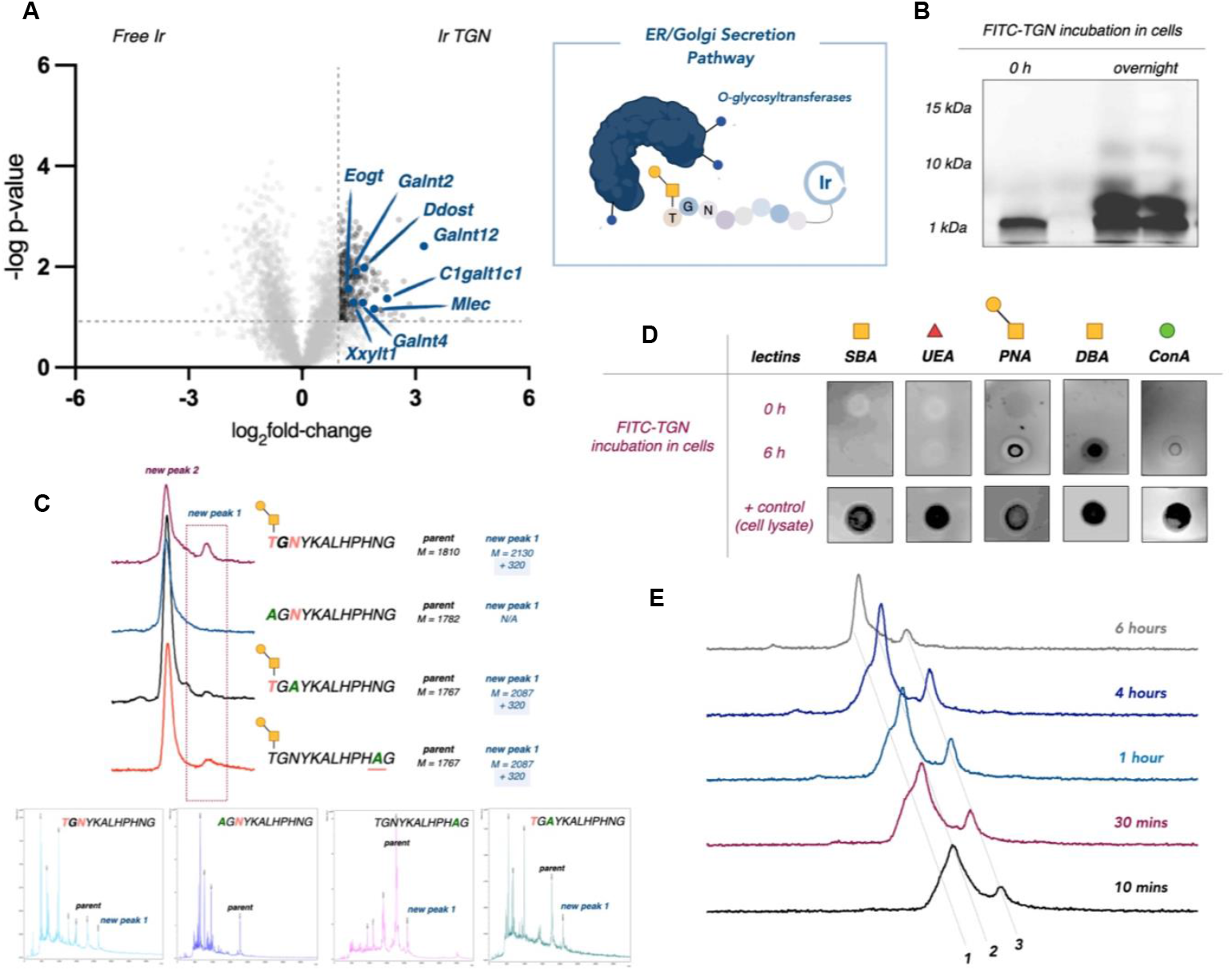
PepTGN is glycosylated during transcytosis. **A.** µMap proteomic analysis of PepTGN in the bEND.3 cell line identifies several O-glycosyltransferases (blue), suggesting that PepTGN may get glycosylated during transcytosis. **B**. Gel electrophoresis of fluorescein labelled PepTGN after incubation with cells displayed multiple new products. **C**. PepTGN analogs were synthesized, replacing potential O- or N-glycosylation sites (Thr-12, Asn-10, Asn-2), identifying Thr-12 as a key residue for glycosylation. **D**. To further confirm glycosylation, we used a biotinylated lectin dot-blot, which showed positive binding for DBA and PNA lectins in PepTGN cellular samples, indicating O-linked glycans. **E**. UPLC-MS analysis of the purified peptides samples revealed a cleavage fragment and a second product with a +322 Da mass shift, suggesting a glycosylation event (1 = cleavage product, 2 = parent peptide & 3 = glycosylated product).

Characterization of the isolated peptides using Ultra Performance Liquid Chromatography-Mass Spectrometry (UPLC-MS) (Figure 6E) revealed one of the new products to be a cleavage fragment of the peptide. However, the second peak also showed a mass shift of +322 Da, which could indicate a glycan modification, possibly involving two distinct glycan attachments.

To further examine this hypothesis, we employed a lectin dot-blot assay to assess the PepTGN glycosylation status before and after incubation with cells (Figure 6D). We immobilized the peptides on a nitrocellulose membrane and observed positive binding in treated samples with the dolichus biflorus agglutinin (DBA) lectin, which recognizes non-terminal *N*-acetylgalactosamine (GalNAc). Similar results were observed with peanut agglutinin (PNA), which recognizes Gal-β(1-3)-GalNAc disaccharides. The PNA lectin is selective for O-linked glycans, suggesting the lone Thr residue in PepTGN as the likely site of modification. To explore this hypothesis, we synthesized a series of PepTGN analogs bearing mutations at each potential site of O- or N-glycosylation: Thr-12, Asn-10, and Asn-2. These analogs were incubated with BBB cells and assessed *via* UPLC-MS (Figure 6C). Interestingly, only the T12A analog did not display a +322 mass shift, indicating that Thr-12 is the likely site of glycosylation.

We believe that this demonstrates how the μMap platform can be used to identify cellular trafficking mechanisms across the BBB that could be useful knowledge to inform drug delivery to the brain. Future work will harness and apply the knowledge of the PepTGN entry receptor (*CALCRL*) and trafficking mechanisms to non-BBB permeable drugs, improving their BBB penetration efficiency.

## Conclusions

We describe herein the development and validation of a µMap photoproximity labeling platform to study the cellular trafficking mechanisms of various CPPs. Using HIV-1 TAT as a proof-of-concept, we demonstrated the ability to map peptide localization at different time points, capturing both known cell-surface proteoglycan interactors and nuclear proteins at the final destination of the peptide within the cell. Building on this demonstration, we next explored CPPs with unidentified cellular entry mechanisms, focusing on two peptides with distinct ionic properties. Through µMap, microscopy, and binding assays, we identified their respective entry receptors, highlighting the diversity in their interaction profiles.

Finally, we sought to map the transcytosis pathway of PepTGN, a BBB-permeable peptide identified *via in vivo* phage display, to gain insight into its elusive intracellular trafficking mechanisms. Time-gated μMap experiments in two BBB-model cell lines allowed us to identify a potential entry receptor (*CALCRL*) and to delineate intracellular trafficking mechanisms and stages through modulation of the peptide’s incubation time. These studies suggested that the peptide likely undergoes glycosylation upon cellular internalization, and further experiments supported this hypothesis and revealed a specific Thr residue within the BBB-targeting peptide sequence as the likely site of glycan attachment.

In conclusion, the μMap platform is well-suited for identifying biomolecular transit routes, revealing novel BBB entry receptors, and elucidating mechanisms that could be targeted for enhanced tissue delivery. Additionally, this technology offers significant potential as a powerful tool for identifying interacting partners of peptides discovered through HTS methods, such as phage or mRNA display. Together, the findings enhance our understanding of peptide trafficking and provide a foundation for improving drug delivery systems to the brain. Future work will focus on leveraging these insights to optimize strategies for targeting the brain and other tissues more effectively.

## Acknowledgements

Research reported in this work was supported by the National Institute of General Medical Sciences of the National Institutes of Health (R35GM134897), the NIH S10 award 1 (S10OD028592-01A1), the Princeton Catalysis Initiative, and kind gifts from Merck, Pfizer, Janssen, Bristol Myers Squibb, Genentech, and Genmab. The authors thank Gary Laevsky and Sha Wang at the Princeton Confocal Proteomics Facility, and Christina DeCoste and Gabriel Palmiere at the Princeton Flow Cytometry Facility. The authors would like to thank Rebecca Lambert for help with editing and submission. Generalized schemes were created using Bio-render.

## Author Contributions

D.C.M. and D.W.C.M. conceived the work. All experiments were performed by D.C.M., S.D.K. and C.P. D.C.M. and D.W.C.M. wrote the manuscript with input from all authors.

